# Inferring intrinsic and extrinsic noise from a dual fluorescent reporter

**DOI:** 10.1101/049486

**Authors:** Erik van Nimwegen

## Abstract

Dual fluorescent reporter constructs, which measure gene expression from two identical promoters within the same cell, allow total gene expression noise to be decomposed into an extrinsic component, roughly associated with cell-to-cell fluctuations in cellular component concentrations, and intrinsic noise, roughly associated with inherent stochasticity of the biochemical reactions involved in gene expression [1]. A recent paper by Fu and Pachter presented frequentist statistical estimators for intrinsic and extrinsic noise using data from dual reporters [2]. For comparison, I here present results of a Bayesian analysis of this problem. I show that the orthodox estimators suffer from pathologies such as predicting negative values for a manifestly non-negative quantity, i.e. variance, and show that the Bayesian estimators do not suffer from such pathologies. In addition, I show that the Bayesian analysis automatically identifies that optimal estimates of intrinsic and extrinsic noise depend on a subtle combination of two statistics of the data, allowing for accuracies that are up to twice the accuracy of the orthodox estimators in some parameter regimes.

I hope up this little worked out example contrasting orthodox statistical analysis based on ad hoc estimators with estimators resulting from a Bayesian analysis, will be educational for others in the field. I distribute a Mathematica Notebook with this paper that allows users to easily reproduce all results and figures of the paper.

## Introduction

How much of the variation in the expression of a gene across a set of single cells is caused by variations in the internal cellular environment of each cell, such as the concentrations of various molecules that are involved in gene expression, and how much variation would exist even if every cell had the exact same cellular environment? To answer this question, a seminal early paper in the field of stochastic gene expression by Elowitz, Swain, and Siggia [1], introduced a dual reporter construct in which two different fluorescent proteins that are driven by identical promoter sequences are integrated into the genome. Since, within each cell, each of the two promoters has experienced an identical cellular environment, the difference in expression of the two reporters within a single cell quantifies the extent to which gene expression varies, even within a constant environment. The authors defined this variability of the reporters within single cells as the ‘intrinsic’ noise, and used this to decompose the total variance into this intrinsic component, and the rest, which they called ‘extrinsic noise’. To illustrate the basic idea, let (*x*_*i*_, *y*_*i*_) be the expression levels (i.e. fluorescence) of the first and second reporter in cell *i* and assume we have measured such pairs of expression levels in *n* cells. The observed mean expression levels are

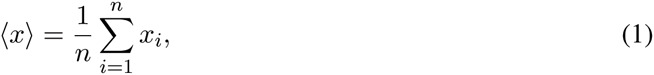

and

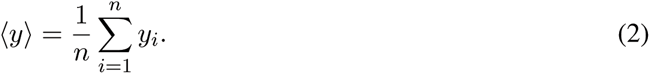

The total amount of observed variance of the two reporters across the *n* cells is

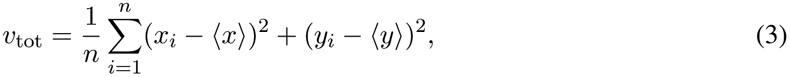

If we define the intrinsic variance *v*_*i*_ as half of the average squared difference of the two reporters in each cell (half because each of the two promoters is subject to this noise)

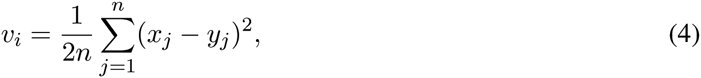

then it is easy to see that the total variance can be decomposed as

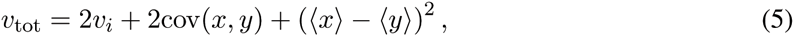

where the covariance of the sample is given by

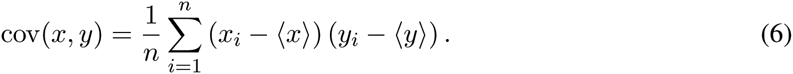

In [1] this covariance is defined to be the extrinsic variance, i.e.

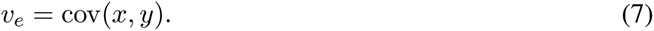

When the two reporters have identical means, or when the data are normalized such that each reporter has equal mean by construction, then the last term in equation (5) is zero, and the total variance vtot naturally decomposes into a sum of the intrinsic variance *v*_*i*_ and an extrinsic variance *v*_*e*_, i.e. *v*_tot_ = 2*v*_*i*_ + *v*_*e*_. Finally, the intrinsic and extrinsic *noise* levels are defined by dividing the variances by the respective mean-squared, i.e. 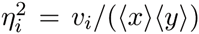 and 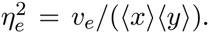. Again, in [1] it is essentially assumed that the means are equal, i.e. 〈*x*〉 = 〈*y*〉.

So far the intrinsic and extrinsic variances where defined directly in terms of the data, i.e. without referring to an underlying probabilistic model. In a recent paper by Fu and Pachter [2], a more explicit probabilistic model was formulated for the dual reporter system, and statistical estimators were constructed, both unbiased estimators, as well as estimators that minimize mean squared error. At a recent visit to Berkeley, Lior brought this work to my attention and invited me to comment on it. In reply I promised to calculate the Bayesian solution for this problem, and to discuss further if this Bayesian solution turns out to differ substantially from the orthodox statistical estimators.

In my opinion, this little problem turns out to be a highly illustrative and educational example of why the Bayesian approach is so superior to an approach using estimators. It is known that, when the estimators are not sufficient statistics, they can exhibit pathological behavior, see e.g. [3, 4] for concrete examples. In this particular example, the orthodox estimators can predict negative values for a variance. Second, as we will see below, while the orthodox estimators per definition use a single statistic of the data, the Bayesian analysis shows that optimal estimates depend on a non-trivial combination of two different statistics of the data.

Below I first introduce the basic setup of the problem and derive the form of the likelihood function. I then discuss the maximum likelihood (ML) solution, which is most similar to the solutions obtained with orthodox estimators. Analysis of the ML solution already clarifies why the orthodox estimators can give absurd estimates in certain parameter regimes. I then derive Bayesian marginal posterior distributions for the intrinsic and extrinsic noise and discuss their intricate dependence on two different statistics of the data. In particular, I will discuss how the Bayesian analysis automatically incorporates subtle dependencies in the inference of the intrinsic and extrinsic noise levels that are not captured by the orthodox estimators.

## Set-up, assumptions, and likelihood

We assume that there are *n* measurements, from *n* cells, of fluorescent reporter expression levels (*x*_*i*_, *y*_*i*_), where 1 ≤ *i* ≤ *n* is the index of the cell, *x*_*i*_ is the fluorescence of the first reporter and *y*_*i*_ is the fluorescence of the other reporter in the cell. In the paper of Elowitz et al. [1] these are CFP and YFP proteins but the setup in principle applies to any dual reporter system.

To construct a likelihood model we make essentially the same assumptions as Fu and Pachter [2]. We assume the two reporters are constructed such that, within the same cell, the values *x*_*i*_ and *y*_*i*_ are drawn independently from the same distribution that has some mean *μ*_*i*_ and variance *v*_*i*_. We will make the assumption that this distribution is a Gaussian

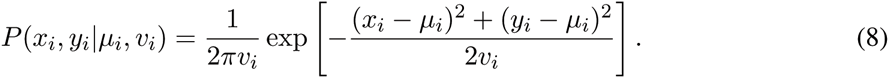

As an aside, since the Gaussian is the maximum entropy distribution conditioned on a given mean and variance, assuming the Gaussian form arguably corresponds to the most general analysis conditioned on these statistics only. As a second aside, we have here assumed that *x* and *y* come from an identical distribution. We could relax this and assume that the means of *x* and *y* differ by a constant amount or by a multiplicative factor, but since this analysis is mainly for illustrative purposes, we will not pursue this more general formulation.

For each cell *i*, the mean *μ*_*i*_ and variance *v*_*i*_ are themselves assumed to be drawn from some distribution as well. Note that *v*_*i*_ is effectively what is meant with the intrinsic noise for cell *i*. Thus, if *v*_*i*_ were to significantly vary across cells, we would have to speak about a *distribution* of intrinsic noise values rather than a given intrinsic noise. To simplify the problem, we follow Fu and Pachter [2] and assume that *v*_*i*_ = *v in every cell*. That way, there is an unambiguous intrinsic noise *v*. At this point it is good to note that our basic biophysical understanding suggests that there is at least a Poisson component to the intrinsic noise level in cell *i* so that the variance *v*_*i*_ must be at least as large as *μ*_*i*_, i.e. *v*_*i*_ ≥ *μ*_*i*_, and this suggests that intrinsic noise levels will likely scale with the mean expression level *μ*_*i*_ of each cell. Thus, in a more serious analysis of this problem we would want to take such information into account. We likely would also want to take into account that the measurements *x*_*i*_ and *y*_*i*_ are also subject to measurement noise, and maybe also that genealogically close cells, such as sister cells that derived from a cell division, may show non-negligible correlation in *μ*_*i*_, and so on. However, since our aims here are mainly to use this problem for illustrative purposes, we will ignore all these more subtle issues.

For the means *μ*_*i*_ we will assume that these have a distribution with overall mean *μ* and variance *v*_*μ*_. Again, the max-ent assumption is to let this distribution be Gaussian as well:

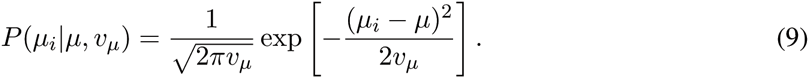

With these assumptions, the probability to obtain values (*x*_*i*_, *y*_*i*_) from a single cell is given by marginalizing over the unknown mean *μ*_*i*_ of the cell *i* in question:

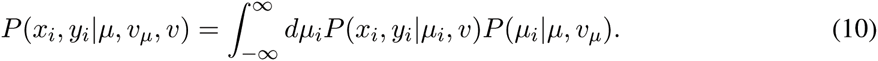

This integral can be easily performed by completing squares in the exponent and yields:

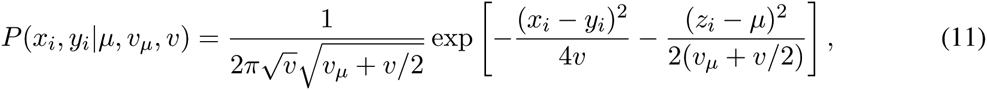

where I have defined *z*_*i*_ = (*x*_*i*_ + *y*_*i*_)/2 to be the average of the two reporters in cell *i*.Note that the result (11) makes intuitive sense. The difference between *x* and *y* is Gaussian distributed with a variance that is the sum of the intrinsic variances of the two reporters, i.e. *2v*. The difference between *μ* and the average of the two reporters is Gaussian distributed with variance *v*_*μ*_ + *v*/2, where the last term in the variance comes from the standard-error of the deviation of the mean of the two reporters from the ‘true mean’ *μ*_*i*_.

Since each cell is an independent measurement from this distribution, the probability of the entire data-set is given by

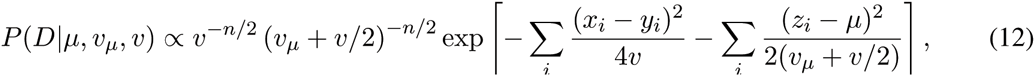

where I left out some constant prefactors that depend on *n* and *π* which are irrelevant for the final result. Since we do not care about the mean *μ*, we will marginalize over this too. Using a uniform prior over *μ* and integrating it out of the likelihood we obtain

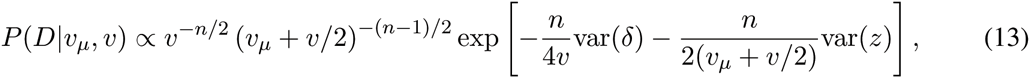

where I have *defined* the empirically observed variances var(*δ*) = Σ_*i*_(*x*_*i*_ − *y*_*i*_)^2^/*n*, var(*z*) = Σ_*i*_(*z*_*i*_ − ⟨*z*⟩)^2^/*n*, and average ⟨*z*⟩ = Σ_*i*_ *z*_*i*_/*n*.

Equation (13) shows that, besides the number of data points *n*, inferences about *v* and *v*_*μ*_ depend on the data only through the statistics var(*δ*) and var(*z*). However, (13) also shows that these two statistics do not provide information about the parameters *v* and *v*_*μ*_ in an independent manner. The likelihood roughly corresponds to a product of two distributions. The first factor *v*^−*n*/2^*e*^−*n*var(*δ*)/(4*v*)^ relates *v* to the statistic var(*δ*) and roughly says that *v* ≈ var(*δ*)/2, with deviations that are proportional to 1/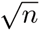. The second factor (*v*_*μ*_ + *v*/2)^−(*n*−1)/2^ e^−*n*var(*z*)/(2*v*_*μ*_+*v*)^ relates the statistic var(*z*) to the sum *v*_*μ*_ + *v*/2, and roughly implies that var(*z*) ≈ *v*_*μ*_ + *v*/2 with deviations proportional to 1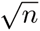. This already makes clear that our estimates of *v* and *v*_*μ*_> will depend on *both* var(*δ*) and var*z*) in a nontrivial manner. However, before I go on with the Bayesian solution to providing estimates for *v* and *v*_*μ*_, we will first derive the maximum likelihood estimates, because these are closest to the orthodox estimators presented in the paper of Fu and Pachter [2].

## Maximum likelihood solution

To obtain the maximum likelihood (ML) estimates for *v* and *v*_*μ*_ we solve for *v*_*_ and *v*_*μ*_*__ such that the derivatives of *P*(*D|v*, *v*_*μ*_) vanish at (*v*_*_, *v*_*μ*_*__). We then obtain

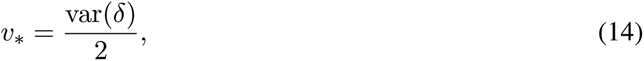

and

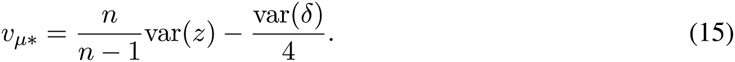

The first of these two is highly intuitive and corresponds closely to the estimators found in the Elowitz et al. paper [1] and in the paper of Fu and Pachter [2], i.e. differing only by terms of order 1/*n* from those estimators. The second expression can be understood as follows. As we noted above, in the limit of large *n*, the statistic var(*z*) will converge to *v*_*μ*_ + *v*/2 and var(*δ*) will converge to 2*v*. Thus, for large *n*, the difference between var(*z*) and var(*δ*)/4 should converge to *v*_*μ*_. As an aside, the reason the ML solution has *n*var(*z*)/(*n* − 1) rather than simply var(*z*) is because one degree of freedom was integrated out of the likelihood (i.e. *μ*).

If we expand var(*z*) and var(*δ*) in terms of the empirical means, variances, and covariance of the (*x, y*) measurements we obtain

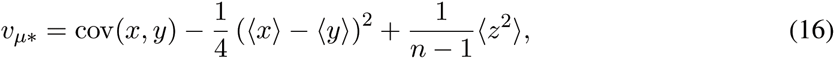

where I have again determined empirical variances and covariances: ⟨*x*⟩ = Σ_*i*_ *x*_*i*_/*n*, ⟨*y*⟩ = Σ_*i*_ *y*_*i*_/*n*, ⟨*z*^2^⟩= Σ_*i*_(*x*_*i*_+*y*_*i*_)^2^/(4*n*), and cov(*x,y*) = −⟨*x*⟩⟨*y*⟩+Σ_*i*_*x*_*i*_*y*_*i*_/*n*. Since *x* and *y* are drawn by definition from distributions with the same mean, their empirical averages over *n* samples will deviate by an amount that scales as 1/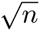. Consequently, both the second and third term in equation (16) are of order 1/*n*. Thus, up to terms of order 1/*n*, the ML estimate is given by the empirical covariance of *x* and *y*, which is precisely the way in which extrinsic variance was defined in [1].

## Orthodox estimators of extrinsic noise can be negative

The estimators for the extrinsic variance presented by Fu and Pachter [2] are also directly proportional to cov(*x*, *y*). For example, the unbiased estimator of *v*_*μ*_ is given by *n*cov(*x, y*)/(*n* − 1). It would thus seem that the ML solution, the estimators of Fu and Pachter, and the original expressions given in [1] all closely agree.

However, it should also be fairly obvious that using a sample covariance as an estimator of a variance is highly problematic. There is no guarantee that the sample covariance will be a positive number! Especially when the true extrinsic variance *v*_*μ*_ is small compared to the intrinsic variance *v*, one can easily get that the covariance of the sample is negative. The estimators would then predict a *negative* value of the extrinsic noise, and that is of course absurd. Note that, depending on the relative size of *v*_*μ*_ and *v*, this may in fact be a very common occurrence. In particular, when the extrinsic variance is negligible compared to the intrinsic variance, the sample covariance will come out negative approximately half of the time.

To illustrate this, I created a data-set by setting *v* = 1, and *v*_*μ*_ = 10^−4^, i.e. with extrinsic fluctuations being 100-fold smaller than the intrinsic fluctuations, and sampled *n* = 25 data points. The first two times I randomly sampled such a synthetic dataset, I obtained positive covariance, but the third time I obtained a dataset with negative covariance. This dataset is shown in the top left panel of Fig. 1. For this dataset, we have var(*δ*) ≈ 2.14, which is only slightly larger than the value of 2*v* = 2 which is expected based on the true value *v* = 1. Similarly, the observed value of var(*z*) ≈ 0.36, which is smaller than the expected *v*_*μ*_ + *v*/2 = 0.5001, but such fluctuations can of course occur with finite datasets. Importantly, the sample covariance comes out negative, i.e. cov(*x, y*) ≈ −0.23, and the estimators would thus estimate a negative value for *v*_*μ*_.

**Figure 1:**
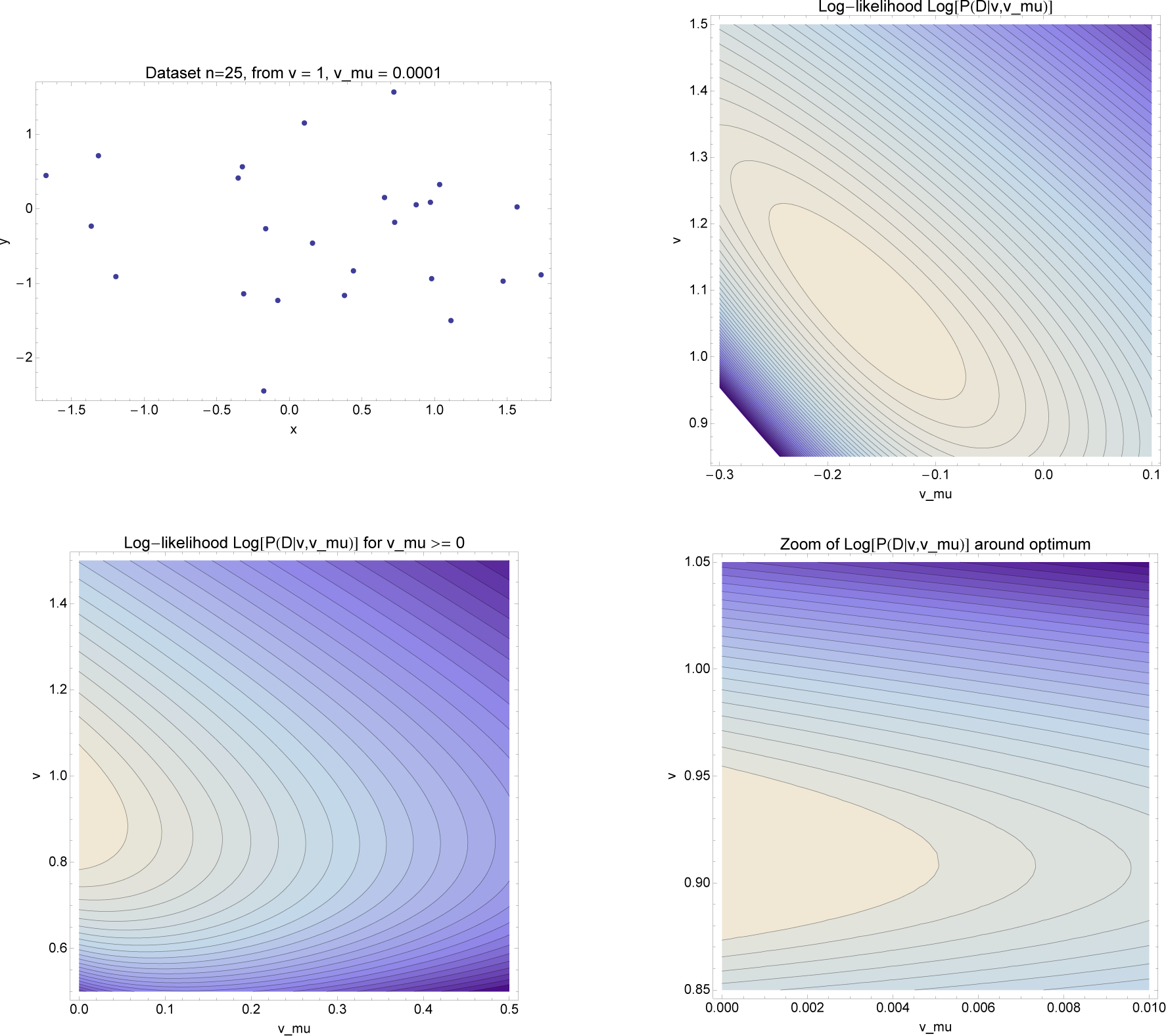
Inference of (*v*, *v*_*μ*_) for a dataset with negative covariance. **Top left:** The *n* = 25 data-points sampled according to the model with *v* = 1 and *v*_*μ*_ = 0.0001. For this particular data-set the relevant empirical statistics of the sample are var(*δ*) = 2.14, var(*z*) = 0.36, and cov(*x,y*) = −0.23. **Top right:** Contourplot of the log-likelihood *log*[*P*(*D|v, v*_*μ*_)] including unphysical values at *v*_*μ*_ < 0. The maximum of the likelihood surface occurs at (*v*_*_ = 1.07, *v*_*μ*\*_ = −0.16). **Bottom left:** The log-likelihood surface restricted to *v*_*μ*_ ≥ 0. **Bottom right:** Close-up of the log-likelihood surface around the optimum (*v*_*_ = 0.91, *v*_*μ*\*_ = 0) when restricted to *v*_*μ*_ ≥ 0.

## Correct maximum likelihood solution

One may ask whether this problem also affects the maximum likelihood estimate. The answer is: it depends on whether one is careful when applying maximum likelihood. Above we obtained the maximum likelihood estimates of *v* and *v*_*μ*_ by differentiating (13) with respect to *v* and *v*_*μ*_ and demanding that these derivatives are zero. However, there is no guarantee that the maximum of the likelihood lies in the physically meaningful region *v* ≥ 0, *v*_*μ*_ ≥ 0. From equation (16) we see that when var(*δ*) > 4*n*var(*z*)/(*n* - 1), the maximum of the likelihood function lies at a *negative* value of *v*_*μ*_. Indeed, for the dataset of Fig. 1, the optimum of the likelihood occurs at a negative value of *v*_*μ*_, as illustrated in the top-right panel of Fig. 1. The naive maximum-likelihood estimate of equations (14) and (16) give *v*_*_ ≈ 1.07, and *v*_*μ*_*__ ≈ −0.16, which is equally absurd.

Of course, the correct way of applying maximum likelihood is to make use of the information that *v* and *v*_*μ*_ are non-negative quantities and find the maximum of the likelihood on the domain *v* ≥ 0, *v*_*μ*_ ≥ 0. One then finds that, whenever var(*δ*)/4 > *n*var(*z*)/(*n* - 1), the correct maximum is at

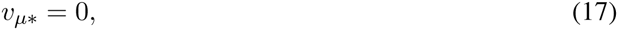

and

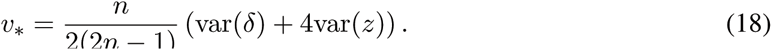

For the data-set of Fig. 1, this optimum occurs at *v*_*_ ≈ 0.91 and *v*_*μ*_*__ = 0.

## Bayesian posteriors for *v*and *v*_*μ*_

In the Bayesian analysis, the next step after obtaining the likelihood is to obtain a joint posterior distribution *P*(*v,v*_*μ*_,*|D*) by multiplying the likelihood *P*(*D|v, v*_*μ*_) by a prior *P*(*v, v*_*μ*_) that represents our prior information about *v* and *v*_*μ*_ (which for example includes that these are non-negative quantities), and normalizing, i.e.

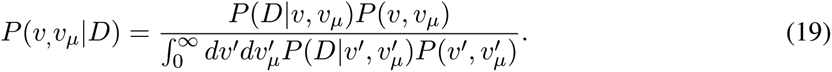

For simplicity, we will assume a uniform prior, which gives *P*(*v, v*_*μ*_*|D*) ∝ *P*(*D|v, v*_*μ*_). From the point of view of the Bayesian analysis, the posterior represents all the information that the data provides about *v* and *v*_*μ*_. Thus, for any region *R* in the upper-quadrant *v* ≥ 0, *v*_*μ*_ ≥ 0, we can calculate the probability that the true values of *v* and *v*_*μ*_ lie in *R* by integrating the posterior over this region.

It is natural to be interested in giving credible intervals for *v* and *v*_*μ*_ separately, and the next logical step in the Bayesian analysis is thus to calculate marginal posterior distributions *P*(*v|D*) and *P*(*v*_*μ*_*|D*). With these, one can then determine, for example, symmetric 90% credible intervals [*v*_min_, *v*_max_] such that *P*(*v* < *v*_min_*|D*) = 0.05 and *P*(*v* ≥ *v*_max_*|D*) = 0.05, and similarly for *v*_*μ*_. Such intervals would give estimates of the ranges of values in *v* and *v*_*μ*_ that are consistent with the data *D* (when considering *v* and *v*_*μ*_, separately).

The marginal posterior distributions for *v* and *v*_*μ*_ are given by the integrals

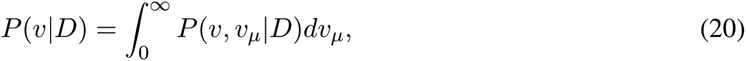

and

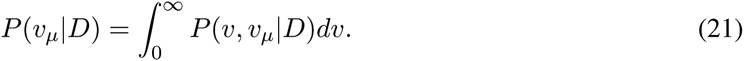

In the Bayesian approach, these posterior distributions contain all information about the values of *v* and *v*_*μ*_ In order to make the question for a single ‘best’ estimate of *v* or *v*_*μ*_ well-defined, we must specify how costly different possible errors in this ‘best estimate’ would be. That is, we need to specify a loss or cost function *C*(*v*^*e*^, *v*^*t*^) that calculates the loss or cost for estimating *v*^*e*^ when the true value of *v* is *v*^*t*^. One can show that the optimal estimate *v^e^* is the one that minimizes the expected loss

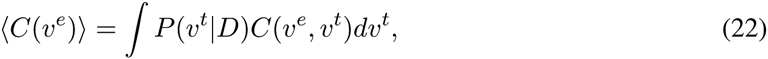

where the expectation is taken with respect to the posterior distribution *P*(*v|D*) [4]. Commonly used loss functions are the squared-deviation *C*(*v*^*t*^, *v*^*e*^) - (*v*^*t*^, *v*^*e*^)^2^ and absolute deviation *C*(*v*^*t*^, *v*^*e*^) = |*v*^*t*^ − *v*^*e*^|. These lead to optimal estimators that are given by the expected mean

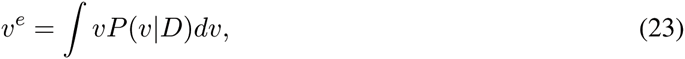

and median

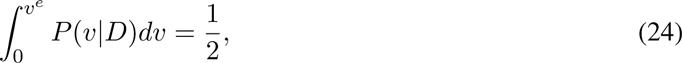

respectively.

These estimators *v*^*e*^ are optimal in the following sense. Imagine the following game played many times: We pick values of *v* and *v*_*μ*_ from the prior distribution *P*(*v, v*_*μ*_) (for example uniform over some range) and generate a data-set *D* of *n* data-points using the model. We then calculate the posterior marginal distributions *P*(*v|D*) and *P*(*v*_*μ*_*|D*) and use these to calculate either the expected means (when minimizing expected square deviation) or expected medians (when minimizing expected absolute deviation). In the limit of many repetitions of this game, the total loss (summed over all repetitions) will be minimized by this procedure, and it will outperform all other estimators in this sense.

## Marginal posterior for intrinsic noise *v*

The integral (20) can be performed analytically and we obtain for the posterior of *v*:

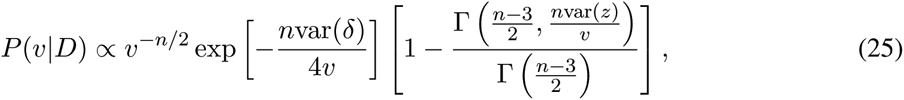

where the incomplete gamma function is defined as

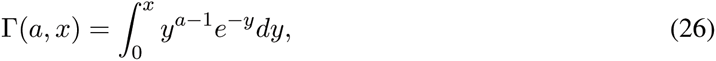

and the ratio 0 ≤ Γ(*A, x*)/Γ(*A*) ≤ 1 is also known as the regularized incomplete gamma function.

It is instructive to analyze this posterior *P*(*v|D*) in some detail. Like the likelihood (13), the posterior naturally decomposes into two factors. The first factor, *v*^−*n*/2^e^−*n*var(*δ*)/(4*v*)^, is the same as the first factor in the likelihood and says, roughly speaking, that *v* must be close to var(*δ*)/2. If the posterior consisted only of this factor, then inferences about the intrinsic noise *v* would only depend on var(*δ*). The maximal posterior value of *v* would be var(*δ*)/2 and the posterior mean (which minimizes expected square loss) would be nvar(*δ*)/(2(n - 4)). These estimates are very similar to the estimator given by Elowitz et al. [1] and the estimators proposed in the Fu and Pachter paper [2]. Thus, these estimators essentially only use the statistic var(*δ*) to estimate *v* and ignore other features of the data.

However, this is not the only factor in the Bayesian posterior (25)! There is a second factor (the part with the incomplete gamma-function within square brackets) that depends on the data through var(*z*). To understand the meaning of this factor, Fig. 2 (left panel) shows the full posterior and the two factors out of which it is composed for the data-set with *v =* 1 and *v*_*μ*_ = 0.0001 that I introduced above.

If the posterior consisted only of the first factor *v*^−*n*/2^e^−*n*var(*δ*)/(4*v*)^, it would be given by the red curve in the left panel of Fig. 2. It would take on a maximum at *v*_*_ = var(*δ*)/2 ≈ 1.07, its mean would be ⟨*v*⟩ ≈ 1.27, its median v_med_ ≈ 1.20, and the symmetric 90% posterior probability interval would be[*v*_min_,*v*_max_] = [0.76,2.04].

**Figure 2:**
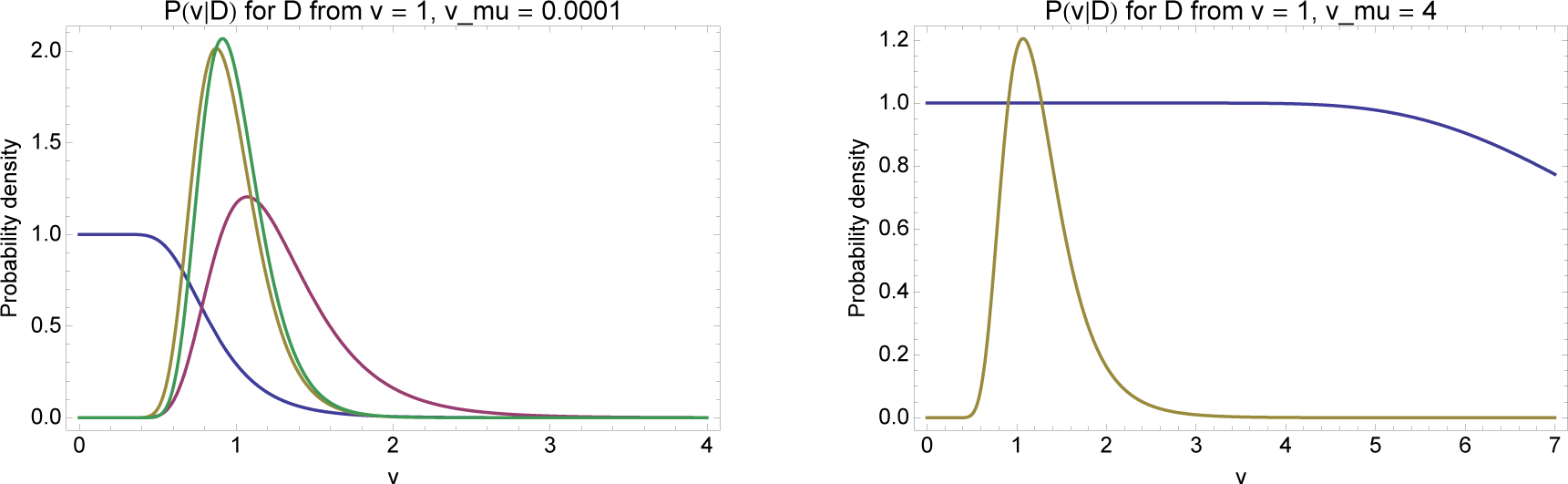
Decomposition of the posterior distribution *P*(*v|D*) for two different data-sets. **Left panel:** The full posterior distribution *P*(*v|D*) (yellow curve) for the data-set of *n* = 25 points sampled from the distribution with *v* = 1 and *v*_*μ*_ = 0.0001 introduced previously. The red curve shows the part of the posterior *v*^−*n*/2^e^−*n*var(*δ*)/(4*v*)^ that depends on var(*δ*) only, whereas the blue curve shows the factor [1 - *Γ*((n - 3)/2,*n*var(*z*)/*v*)/*Γ*((*n* - 3)/2)] that depends on var(*z*). The yellow curve is the full posterior obtained by multiplying the factors represented by the red and blue curves. The green curve shows the posterior *P*(*v|D*, *v*_*μ*_ = 0) that would be obtained if it were known that there is no extrinsic noise, i.e. *v*_*μ*_ = 0. **Right panel:** The same curves (except the green curve) for a data-set of *n* = 25 points sampled from the distribution with *v* = 1 and *v*_*μ*_ = 4.

However, there is also a factor depending on the variance var(*z*) of the average expression *z* = (*x*+*y*)/2 across the cells, shown in blue. This factor is 1 for small values ofv and then starts dropping at some critical value ofv, which is approximately at *v* = 2*n*var(*z*)/(*n* − 3), which is equal to 0.82 for this dataset. Because of this, the full posterior (yellow curve) is shifted toward slightly smaller values of *v*, and it becomes significantly more tightly peaked. The yellow posterior takes on a maximum at *v*_*_ ≈ 0.91, its mean equals ⟨*v*⟩ ≈ 0.96, its median equals *v*_med_ ≈ 0.93 and its symmetric 90% posterior probability interval equals [*v*_min_, *v*_max_] = [0.65,1.36]. Importantly, the 90% posterior probability interval is almost half as wide for the full posterior (yellow curve) compared to posterior based on the first factor only (red curve). That is, for this data-set, inclusion of the second factor that depends on the statistic var(*z*) almost *doubles the accuracy* of the estimate.

Intuitively, the blue curve represents the impact on *v* of information about *v*_*μ*_, which is encoded in the statistic var(*z*). So how can information about *v*_*μ*_ lead to an almost doubling of the estimate of *v*? To understand this, let’s imagine we *knew in advance* that *v*_*μ*_ is very small. For illustration, let’s assume that we *know v*_*μ*_ = 0, i.e. that there is no extrinsic noise at all. In that case the *x* and *y* measurements in each cell become independent. That is, in each cell *i* the measurements *x*_*i*_ and *y*_*i*_ are drawn from independent Gaussians with mean *μ* and variance *v* and the likelihood becomes

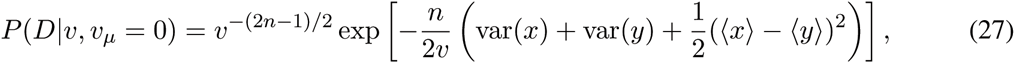

which has a maximum at

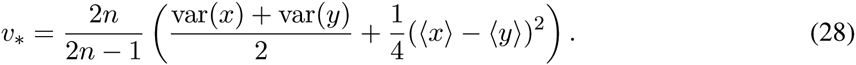

Up to corrections of order 1/*n*, this is equal to the *average* of the variances var(*x*) and var(*y*). Thus, when we know *v*_*μ*_ = 0, then we base our estimate for *v* on *2n* independent measurements: *n* measurements determining the variance var(*x*) and *n* separate measurements determining the variance var(*y*). In contrast, with the red curve *v*^−*n*/2^e^−*n*var(*δ*)/(4*v*)^, we are basing our estimate ofv on only *n* measurements of the difference (*x − y*) in each cell. This explains why, if we know *v*_*μ*_ = 0, then the accuracy of our estimate of *v* can become much larger.

This should elucidate that the effect of incorporating knowledge of var(*z*), i.e. the effect of the blue curve in Fig. 2, is similar to *knowing* that *v*_*μ*_ = 0. To confirm this, the green curve in the left panel of Fig. 2 shows the posterior *P*(*v|D, v*_*μ*_ = 0) that would be obtained if it were known that *v*_*μ*_ = 0. Indeed, we see that the full posterior (yellow) curve, is quite close to the posterior that would be obtained if we knew *v*_*μ*_ = 0. Thus, by incorporating the information var(*z*) contains, we have effectively incorporated knowledge that *v*_*μ*_ is very small.

Let’s now see how the blue accomplishes this. Remember that the likelihood (13) naturally decomposed into afactor *v*^−*n*/2^e^−*n*var(*δ*)/(4*v*)^ (the red curve) andafactor (*v*_*μ*_+*v*/2)^-(*n*-1)/2^e^−*n*var(*z*)/(2(*v*_*μ*_+*v*/2))^. This second factor, roughly speaking, says that *v*_*μ*_+*v*/2 should be close to var(*z*). The blue curve results from integrating this factor over *v*_*μ*_. Again speaking very roughly, this integral measures how many ways *v*_*μ*_ can be set to match the requirement that (*v*_*μ*_ + *v*/2) is close to var(*z*). If *v*/2 is small compared to var(*z*), one can always match this requirement by picking a large value of *v*_*μ*_ and so the contribution of the integral is approximately constant. However, when *v*/2 becomes close to or even larger than var(*z*), then there is essentially no value of *v*_*μ*_ left that is consistent with the observed var(*z*), i.e. even with *v*_*μ*_ = 0 the sum *v*_*μ*_ + *v*/2 is already larger than the observed var(*z*), and the integral will become small. In other words, the blue curve roughly constrains *v* to be less than 2var(*z*). At the same time, the red curve constraints *v* to be close to var(*δ*)/2. For our data, these two constraints together are equivalent to knowing that *v*_*μ*_ must be small and, consequently, the full posterior is similar to what would be obtained if one knew that *v*_*μ*_ is small.

Note also that, when *v*_*μ*_ is large, the constraint that *v* < 2var(*z*) will become irrelevant. An example is shown in the right figure of panel Fig. 2. Here I generated a data-set with *v* = 1 and *v*_*μ*_ = 4. Because the red curve depends only on var(*δ*), and var(*δ*) depends only on *v*, and not *v*_*μ*_, the red curve in the right panel is similar to the red curve in the left panel. However, because *v*_*μ*_ is now much larger, var(*z*) is now also much larger, and the blue curve therefore only starts to drop at values of *v* that are outside of the range supported by the red curve. As a consequence,the final posterior *P*(*v|D*) (yellow) is now entirely on top of the red curve *v*^−*n*/2^e^−*n*var(*z*)/(4*v*)^, i.e. the factor represented by the blue curve has no influence in this case.

## Marginal posterior for extrinsic noise *v*_*μ*_

It appears to me that the integral (21) cannot easily be done analytically (although, admittedly, I didn’t try very hard, and I might have missed a clever substitution). However, it is straightforward to perform these integrals numerically to obtain the posterior for *v*_*μ*_. Figure 3 shows the posterior distributions that I obtained numerically for *P*(*v*_*μ*_*|D*) for the same two data-sets. Note that, for the first data-set, *v*_*μ*_ is correctly inferred to be small (at most 0.5 or so) and for the second data-set the maximum of the posterior is close to the true value of *v*_*μ*_ = 4.

**Figure 3:**
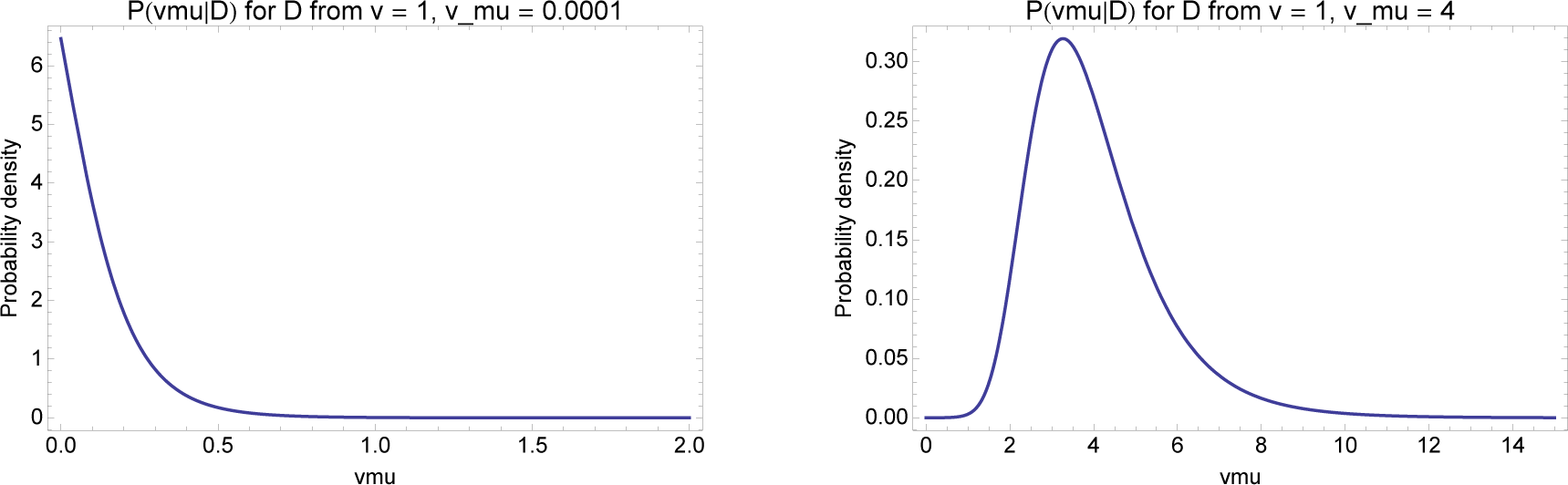
Posterior *P*(*v*_*μ*_*|D*) for the extrinsic noise *vμ* for the same two data-set introduced previously. **Left panel:** The posterior *P*(*v*_*μ*_*|D*) for the data-set with *n* = 25 points sampled from *v* = 1, *v*_*μ*_ = 0.0001. **Right panel:** The posterior *P*(*v*_*μ*_*|D*) for the data-set with *n* = 25 points sampled from *v* = 1,*v*_*μ*_ = 4.

I noticed that if one defines *v*_*μ*_ = λ*v*, i.e. λ = *v*_*μ*_/*v* is the size of the extrinsic noise relative to the intrinsic noise, then the integral over *v* can be performed analytically and one obtains for the posterior of λ:

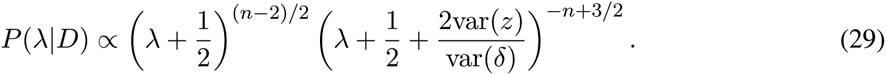

This solution makes clear that the ratio λ depends on the data only through the number of data-points *n* and the ratio

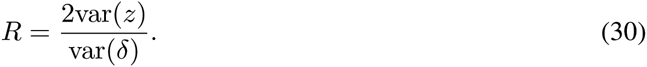

This is another illustration of how a Bayesian analysis can often identify what the relevant statistics of the data are, i.e. here the ratio *R* is the only property of the data that is relevant for inferences regarding λ.

The maximal posterior estimate of λ is given by

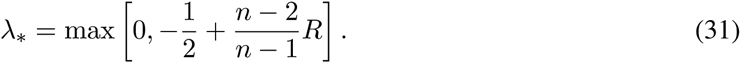

To determine the mean, median and 90% posterior intervals, it is helpful to introduce the parameter

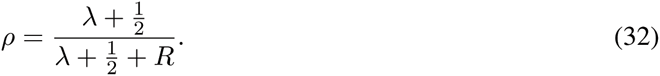

The probability density for *ρ* takes on the form of a beta-distribution:

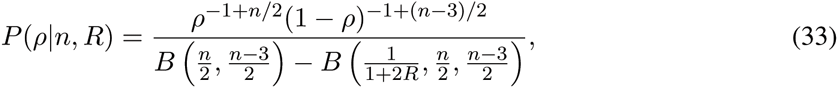

where as λ runs from 0 to ∝, *ρ* runs from *ρ*_min_ = 1/(1 + 2*R*) to 1, *B*(*a, b*) is the beta-function, and *B*(*x, a, b*) is the incomplete beta-function:

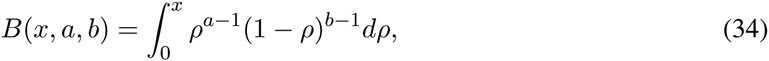

with *B*(*a,b*) = *B*(*1,a,b*). The mean, median and 90% posterior probability interval can then be expressed in terms of incomplete beta-functions. For example we find for the mean

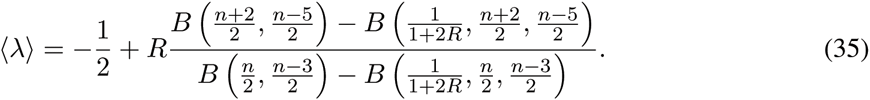

Note that in equation (31), we had to again explicitly check that −1/2 + (*n* − 2)*R*/(*n* − 1) ≥ 0. I leave it as an entertaining exercise for the reader to convince her/himself that, although *R* may well come out smaller than 1/2, through the subtle dependence on *R* of the ratio of beta-functions, the expression (35) is guaranteed to be non-negative.

The median λ_med_ can be obtained by finding *ρ*_med_ that satisfies

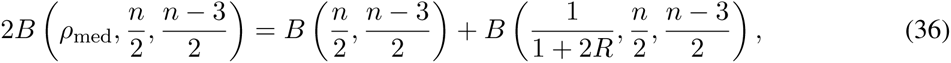

and then setting λ_med_ = −1/2 + *R ρ*_med_/(1 −*ρ*_med_). The symmetric 90% posterior probability interval can similarly be solved in terms of incomplete beta-functions and solutions for this interval are provided in the Mathematica notebook that I distribute with this paper.

To illustrate the behavior of the posterior *P*(λ*|D*), Fig. 4 shows the posterior for the two data-sets with *n* = 25 data-points shown previously, i.e. with *v* = 1,*v*_μ_> = 0.0001 for the first and *v* = 1, *v*_*μ*_ = 4 for the second.

**Figure 4:**
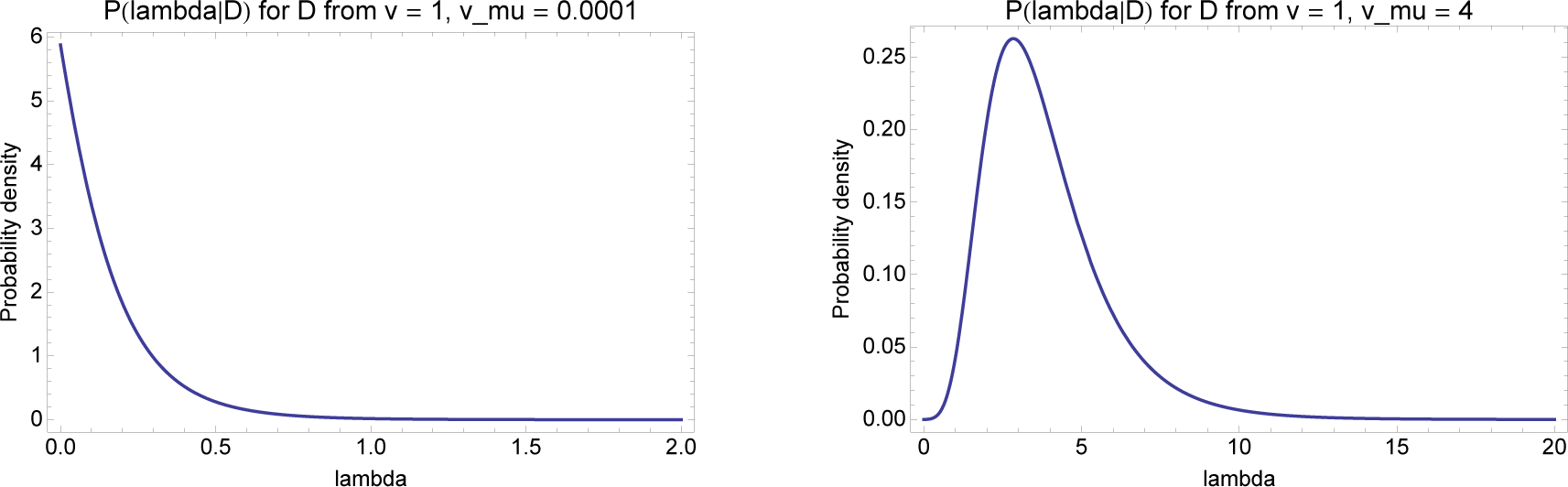
Posteriors *P*(λ*|D*) for the ratio of extrinsic to intrinsic noise λ = *v*_*μ*_/*v*, for the two data-sets. **Left panel:** The posterior *P*(λ*|D*) for the data-set with *n* = 25 points sampled from *v* = 1,*v*_*μ*_ = 0.0001. **Right panel:** The posterior *P*(λ*|D*) for a data-set with *n* = 25 points sampled from *v* = 1, *v*_*μ*_ = 4.

For the first data-set (left panel), the ML value of λ is λ_*_ = 0, the mean is 〈λ〉 ≈ 0.17, the median λ_med_ ≈ 0.11, and the shortest 90% posterior probability interval is [0,0.37]. For the second dataset (right panel) the ML value is λ_*_ ≈ 2.84, the mean 〈λ〉 ≈ 3.85, the median λ_med_ ≈ 3.47 and the symmetric 90% posterior probability interval is [1.49, 7.49]. All these are of course very reasonable given the true values λ = 0.0001 and λ = 4, and the limited amount of data.

## Discussion

I here looked at a simple statistical problem of inferring two unknown parameters (intrinsic and extrinsic variance in gene expression) from measurements using dual reporter constructs. In the approach that I presented, probability theory is seen as an extension of logic [4], and probabilities quantify states of information regarding these unknown quantities. In this approach, the solution to the problem is the calculation of posterior distributions *P*(*v*, *v*_*μ*_*|D*), *P*(*v|D*), and *P*(*v*_*μ*_*|D*) that represent how likely it is that *v* and *v*_*μ*_ have particular values given our data *D*, and any other prior information that we may have had regarding *v* and *v*_*μ*_.

Although the interpretation of probabilities as representing states of information is arguably the original interpretation given by the early developers of the theory such as Bernoulli and Laplace, for most of the 20th century the prevailing view has been that probabilities correspond to frequencies within repeated ‘random experiments’. In this view, probabilities can only be assigned to *random variables*. For this example, the random variables would be the measured fluorescence intensities of the reporters in each cell. In contrast, the parameters *v* and *v*_*μ*_ would just be unknown constants and ‘not random’. Consequently, in this interpretation it makes no sense to talk about ‘the probability that *v* is between this and that value’. Instead, in frequentist statistics one constructs estimators of the unknown quantities that are functions of the data, i.e. in the current problem the estimators of intrinsic and extrinsic variance proposed in [1] and [2] are proportional to var(*δ*) and cov(*x, y*) respectively. The performance of these estimators is then assessed by imagining fixing *v* and *v*_*μ*_, repeatedly sampling measurements from their frequency distributions, calculating the estimators, and then calculating how close these estimators are to the true values on average, e.g. their average squared deviation from the true values. However, note that such performance measures only describe the average performance of the estimator across many experiments. They do not say anything about the particular performance for a single individual data-set with particular var(*δ*) and cov(x, *y*). In fact, it has been shown that if estimators are not based on sufficient statistics, they can show severe pathologies such as predicting values that are known to be impossible [3, 4].

Indeed, our analysis showed that var(*δ*) and cov(*x, y*) are not sufficient statistics for the intrinsic and extrinsic variances, and it is thus not surprising that they indeed show pathological behavior. In the case of cov(*x, y*) the pathology is particularly simple: The estimator can easily estimate a negative value for the extrinsic variance. For the intrinsic variance the situation was more subtle. When *v*_*μ*_ is very small relative to *v*, the estimator var(*δ*) is not producing absurd values, but it is clearly non-optimal,. By comparing the Bayesian estimates with the orthodox estimator we saw that, when *v*_*μ*_ ≈ 0, using var(*δ*) roughly corresponds to throwing away half of the data.

Estimators that can produce absurd values for some datasets, such as cov(*x, y*) for this problem, are sometimes defended on the grounds that, in the particular applications in question, such pathologies are unlikely to occur. Indeed, in the systems studied in [1] the extrinsic variance *v*_*μ*_ was never very small, and the estimator cov(*x, y*) does not exhibit pathological behavior in this case. But this is not a valid defense for the *mathematical procedure* as such. In my opinion, once we know that our method can in principle produce nonsensical results, it is no longer acceptable as a rigorous mathematical procedure. Moreover, as our analysis shows, there is no need to use such estimators. Instead of needing *ad hoc* guesses at the functional form of the estimators, the Bayesian analysis gives posterior distributions for *v* and *v*_*μ*_ that are unambiguous and are easily calculated from first principles. Bayesian estimators that minimize any desired loss function are also easily obtained, e.g. the estimator with minimal expected square deviation is simply the posterior mean, and the estimator with minimal expected absolute deviation is the posterior median.

We studied *P*(*v|D*), the Bayesian posterior for the intrinsic variance, in some detail and found that it automatically incorporates rather subtle dependencies between *v*, *v*_*μ*_, and the two statistics var(*δ*) and var(*z*). In certain parameter regions both var(*δ*) and var(*z*) contain useful information about *v* and the Bayesian posterior then automatically incorporates that information, while in other parameter regimes almost all relevant information is in var(*δ*) and the Bayesian posterior then automatically reduces to a form that effectively only depends on var(*δ*). Importantly, these results did not rely on us inventing clever estimators or developing specific new procedures for this problem. We simply used the standard rules of probability theory to calculate the posteriors *P*(*v|D*) and *P*(*v|D*). This *automatically* incorporated all the subtle dependencies between *v*, *v*_*μ*_ and the data. We in fact had to do quite some interpreting work after the fact to understand how the Bayesian analysis was incorporating these subtle dependencies.

As another example, the Bayesian analysis also showed that, whereas there are no sufficient statistics for either *v* or *v*μ*, the* ratio of sample variances *R* = 2var(*z*)/var(*δ*) is a sufficient statistic for the noise-ratio λ = *v*_*μ*_/*v* and we derived an explicit expression for an estimator with minimal expected quadratic deviation for this ratio in equation (35).

It is interesting to note that the two reporter construct of Elowitz et al. [1] was designed specifically to be able to estimate intrinsic noise from measurements of expression differences *δ*_*i*_ = *x*_*i*_ − *y*_*i*_ across different cells *i*. However, when we performed the Bayesian calculation we found that not only the statistic var(*δ*) carried information about *v*, but that var(*z*) can carry subtle information about *v* as well.

This kind of story by now feels very familiar to me. I encountered Jaynes’ book on probability theory interpreted as an extension of logic [4] during my PhD and I have been applying this methodology in my own research ever since. By now it has happened many times that, when estimating some quantity of interest from experimental data, intuition suggested that all information regarding the quantity of interest should be contained in some single simple statistic of the data, only to find that the proper probability calculation suggested a significantly more complex function of the data. After studying this more complex function in more detail I would find, time and time again, that the Bayesian calculation was automatically incorporating all kinds of subtle dependencies that I had not anticipated beforehand. It is exactly these kinds of experiences that most strongly convinced me that calculations based on treating probability theory as an extension of logic are vastly superior to more orthodox methods that demand one think of probability only as a frequency in repeated experiments. My main motivation for writing up the Bayesian solution for this simple problem in some detail is the hope that it might similarly convince some others of the same: You don’t have to restrict yourself to outdated blunt instruments when extracting information from your experimental data. Probability theory as extended logic provides an integrated rigorous theoretical framework that is unambiguous, pathology-free, and often with demonstrably superior performance.

